# Teaching adipose tissue as an organ system: addressing weight bias and enhancing anatomical understanding through *AdipoAtlas*

**DOI:** 10.64898/2026.07.29.741583

**Authors:** Isabel D.K. Hermsmeyer, Kendrin R. Sonneville, Annie M.S. Patterson, Rowan M. Sherwood, Emily R. Orlikoff, Ormond A. MacDougald, Mary E. Orczykowski

## Abstract

Adipose tissue (body fat) is increasingly recognized as a dynamic organ system essential to human health, with distinct depots that serve specialized biological functions. However, this understanding is not reflected in traditional human anatomy curricula, where adipose tissue is typically introduced only as connective tissue and treated in dissection as an obstacle to be removed rather than a structure worthy of study. Anatomy dissection courses have been identified as a source of negative weight bias in medical students, with students reporting disgust toward adipose tissue and frustration with its removal to visualize course-required anatomical structures. Here, we hypothesize that reframing adipose tissue as a functional organ system within anatomy curricula may improve students’ understanding of human anatomy and mitigate these negative perceptions. To test this, we created a human adipose atlas (AdipoAtlas) by identifying and photographing macroscopic adipose depots in an anatomical donor and organizing these images into an anatomical reference with evidence-based functional descriptions. The atlas was integrated into a human anatomy dissection course, followed by a survey assessing students’ perceptions of adipose tissue and attitudes related to weight bias, with a concurrent non-dissection anatomy course serving as a control. Students in the adipose-inclusive course reported more positive perceptions and improved understanding of adipose tissue as a multifunctional organ system, while responses related to weight-based discrimination and broader attitudes toward body size were similar between groups. These results support our hypothesis that integrating adipose tissue into anatomy education improves anatomical understanding and may reduce negative weight-related perceptions.

## Introduction

Adipose tissue comprises a diverse set of lipid-storing tissues distributed across the human body (see supplemental materials, full list). These tissues are typically classified into three major forms: white adipose tissue (WAT), brown adipose tissue (BAT), and bone marrow adipose tissue (BMAT). WAT makes up the majority of adipose in the human body (Abe, 2021). It is found subcutaneously under the skin, viscerally around the internal organs, and in several specialized locations (i.e. intraarticular, retroorbital) (Zwick et al., 2018). In addition to its active metabolic functions, WAT provides structural support and acts as an active immune and endocrine organ, producing signaling molecules that influence both local and systemic physiology (Emont & Rosen; 2023; Grant & Dixit; 2015; Khan et al., 2019; Loft et al., 2025; Luong et al., 2019). BAT makes up a comparatively smaller portion of adipose tissue in the human body, having a more substantial presence in infants but persists into adulthood in small, discrete depots, particularly in the supraclavicular and paraspinal regions. Brown adipocytes are characterized by multilocular lipid droplets and high mitochondrial density, enabling lipid oxidation for non-shivering thermogenesis, making it an important contributor to thermal regulation (Cypess, 2022; Devlin; 2021). BMAT is another subtype of adipose, located within the marrow cavities of bone, making up roughly 10% of total body adipose tissue in adult humans. It is functionally and transcriptionally distinct from white and brown adipose tissues, and has been shown to support both local blood production (hematopoiesis) and bone homeostasis (Li et al., 2022; Li & Rosen, 2023).

Researchers studying adipose tissue are increasingly emphasizing that adipose tissue is a vital organ system (Cinti, 2001; Cypess, 2022; Emont & Rosen, 2023; Kershaw & Flier; 2004). This is evidenced not only by its dynamic physiological functions, but also by the pathological consequences of adipose deficiency. Specifically, age-related adipose loss is associated with metabolic dysfunction and frailty, rapid and/or excessive adipose loss can impair endocrine and immune regulation, and congenital or acquired lipodystrophies result in severe insulin resistance, dyslipidemia, and ectopic lipid deposition (Kershaw & Flier, 2004; Maung et al., 2025; Tchkonia et al., 2010). Consistent with its role as a vital organ system, adipose tissue is not diffusely distributed but instead organized into distinct depots throughout the human body, regardless of individual adiposity (Zwick et al., 2018). This organization underscores its relevance as a structured component of human anatomy. Despite this growing scientific consensus, adipose tissue has been described as largely absent from anatomical sciences education and anatomical atlases (Cypess, 2022). A recent analysis of widely used anatomy textbooks found that although adipose tissue is often visually present, its inclusion is typically limited or incidental, with low labeling rates and terminology that often frames it as non-functional or pathological rather than as an integrated organ system (Hermsmeyer et al., 2026).

Anatomy dissection has been identified as a source of negative weight bias in medical education, which may shape negative perceptions of higher-weight patients among future clinicians (Goss et al., 2020). Weight bias in healthcare settings is widespread and associated with poorer patient–provider interactions, reduced quality of care, and worse health outcomes, independent of patients’ body mass index (Fruh et al., 2021; Phelan et al., 2015a). In their study, Goss et al. (2020) describe the main source of this negative weight bias to be due to disgust and frustration over having to remove and clean away adipose tissue to visualize the structures outlined in their anatomy dissection course curriculum. Consequently, students assigned to donors with larger bodies had more perceived “work” to do than those assigned to thinner donors. Goss et al. additionally describe that students working on donors with higher adiposity were more likely to characterize them as unhealthy, while thinner donors were described as healthy and ‘in shape,’ despite students having minimal information about donor medical histories. Their findings suggest that the treatment of adipose tissue as an obstacle rather than an integral part of anatomy may contribute to the development of weight bias within anatomy education.

In the present study, we hypothesize that inclusion of adipose tissues within anatomy dissection curricula, rather than treatment as an obstruction to be “cleaned away” to see underlying structures, will help reduce these negative effects and strengthen students’ anatomical understanding. To test this, we developed a comprehensive photographic atlas of human adipose depots (AdipoAtlas) and integrated it into a semester-long anatomy dissection course, comparing student perceptions and attitudes to those of a concurrently taught, non-dissection anatomy course without the intervention. This work explores whether reframing adipose tissue as a legitimate anatomical structure within dissection curricula can inform more accurate anatomical learning and mitigate the development of negative perceptions towards adipose tissue, diverse body sizes and weight bias.

## Methods

### Human anatomical donor dissection

A human anatomical donor (n = 1, female) was obtained from the University of Michigan Anatomical Donations Program (Ann Arbor, MI, USA). The donor was anonymous, with only sex, age, and an abbreviated medical history revealed. The authors state that every effort was made to follow all local and international ethical guidelines and laws that pertain to the use of human anatomical donors in research. All donors were treated in accordance with local and national laws and regulations, and their use in generating multimedia was approved by the University of Michigan and included within the donor consent forms.

The donor was lightly embalmed using a formaldehyde solution through the common carotid artery using standardized techniques. Light embalming was used rather than standard embalming to better preserve adipose tissue integrity, as standard embalming has been observed to cause adipocyte rupture and lipid leakage. After embalming, the donor was stored in a 4 degree Celsius cooler to help maintain preservation. We performed a full dissection of the donor, documenting and photographing each distinct adipose depot with a Sony (Alpha ZV-E10 II).

### AdipoAtlas generation and integration into anatomy dissection course materials

All photographs were minimally post-processed using Adobe Photoshop (Version 27.x) to remove the background of the images and any identifying features, standardize lighting, contrast, and to improve visual clarity without altering anatomical structures. These photographs were curated to create a comprehensive photographic record of macroscopically visible adipose depots in the human body.

This AdipoAtlas was then incorporated into anatomy course materials. Within the course Google Slides laboratory guides, the majority of adipose depots were given a dedicated slide consistent with the standard treatment of other anatomical structures, and photographs of depots were supplemented with a verbal description of their anatomical location, structure, and summarized known physiologic function.

Within the lab guides, students were instructed to identify these depots, but typically not to fully reveal them. Clean removal of the dermis to preserve subcutaneous adipose tissue requires substantial time and effort, which was impractical within the time constraints of typical anatomy dissection course structure. For subcutaneous depots, students were instructed to identify distinct depots during skin and fascia removal, whereas for visceral or specialized adipose, they were instructed to identify depots and associated features prior to any removal needed to expose associated organs or vascular tissues.

AdipoAtlas is available for educational use and can be accessed here: www.adipoatlas.com

### Selection of intervention and non-intervention control groups

To test the efficacy of the adipose-inclusive curriculum in reducing negative perceptions of adipose tissue and weight bias, we selected the fall 2025 offering of Anatomy 503, a semester-long elective dissection course at the University of Michigan for upper-division undergraduate and graduate students pursuing diverse health professions and biomedical careers. This setting provided an opportunity to reach future healthcare providers, researchers, and educators early in their training, while serving as an initial testing context for future adaptation to medical school anatomy courses. As the anatomy 503 course only has one class per year, we were unable to include an anatomy 503 course without the adipose tissue inclusive curriculum intervention for direct comparison. Instead, we used the concurrently running anatomy 403 (ANAT 403) class, a course that covers similar material as anatomy 503 prior to adipose tissue inclusion, but does not include manual dissection and instead consists of learning from plastinated donors. This course effectively functions as a negative control group, as they are similarly learning anatomy but are not performing dissection, the identified source of negative perceptions of adipose tissue and weight bias. The anatomy 503 course received the adipose tissue inclusive curriculum, while the anatomy 403 course did not.

### Survey construction, pilot testing and distribution

The survey was constructed based on the structure and questions contained within Goss et al. (2020). However, the survey was modified to meet the needs of this study by adding questions specific to the course structure and adipose tissues. Two individual items were also included from validated weight bias measures described by Cain et al. (2022). These items were selected from different conceptual domains within the Fat Attitudes Assessment Toolkit (FAAT) framework, specifically, Critical Health (‘Body weight isn’t a reliable indicator of health’) and Activism Orientation (‘We need to take weight-based discrimination as seriously as other forms of discrimination’) based on their relevance to the study. To minimize respondent burden and encourage participation, a full weight bias scale was not administered. The final survey consisted of 16 questions, six of which included free-form text box responses (see supplemental materials).

The survey, along with the overall study structure, was submitted for University of Michigan Institutional Review Board (IRB) review. The study [HUM00279629] was deemed exempt from ongoing IRB review, per the federal exemption category 2(i) and/or 2(ii) at 45 CFR 46.104(d). The survey was implemented and distributed using Qualtrics, with settings configured to ensure respondent anonymity and prevent collection of identifying information. Course enrollment (ANAT 403 or ANAT 503) was collected as a required item to permit course-level comparisons; all remaining questions were optional. As an incentive, following completion of the research survey, participants would then have the option to participate in a gift card raffle.

Following review by coauthors, but prior to distribution to study participants, the survey was pilot tested to assess clarity, interpretability, and alignment with the study aims. Pilot testing was conducted with a separate group of graduate and undergraduate students working in laboratory environments with an anatomical component, including individuals from laboratories that study adipose tissue biology and those without specific adipose-focused research. Results and feedback from pilot participants were used to refine question wording, ensure consistent interpretation across respondents with varying levels of anatomical and adipose-specific expertise, and confirm that the survey items captured perceptions of adipose tissue, anatomical relevance, and attitudinal change as intended. No pilot participants were included in the final study sample.

### Statistical analysis

Statistical analyses were conducted on closed-ended survey items only; open-ended text responses were analyzed separately using qualitative methods. Survey responses were analyzed using regression models appropriate to the measurement scale of each item. Conceptualizations of adipose tissue (Q8, see supplemental materials), which were structured as an ordered progression from harmful/pathological to vital organ system, were analyzed using ordinal logistic regression (proportional odds models) to assess course-related differences in the likelihood of endorsing more positive views. Likert-type agreement items from validated weight-bias measures (Q12–Q13) were also analyzed using ordinal logistic regression to model proportional shifts across ordered response categories without assuming equal intervals between levels of agreement.

Survey items assessing self-reported changes in attitudes following the anatomy lab experience (Q9–Q11) were analyzed using multinomial logistic regression, as response categories lacked a natural ordering. For these models, “No change” served as the reference category, and relative risk ratios (RRRs) were used to compare the likelihood of each response category between courses. All models included course (ANAT 403 vs. ANAT 503) as the primary predictor.

Student performance on adipose-related exam questions was assessed across the semester. For each assessment, questions pertaining to adipose tissue were identified, and performance was summarized as the proportion of adipose-related questions answered correctly (number correct divided by total adipose-related questions for that assessment). Mean performance was calculated for each assessment across students and used to visualize overall trends across the semester.

Statistical significance was assessed at α = 0.05. We conducted all analyses in R (v.4.3.2; R Foundation for Statistical Computing), using the ordinal package for ordinal logistic regression and the nnet package for multinomial logistic regression.

### Qualitative analysis

Open-ended survey responses were analyzed using thematic analysis to identify common patterns in students’ perceptions of adipose tissue and body size following the anatomy laboratory experience. Responses were first reviewed in their entirety to achieve familiarity with the data. A simple initial coding framework was developed based on the study design (e.g., more positive, mixed, more negative), with the understanding that a more specific framework would be generated after responses were analyzed and overarching themes observed.

Responses from both courses were coded using this framework to enable comparison between the intervention (ANAT 503) and comparison (ANAT 403) groups. Codes were refined through repeated review to capture key dimensions of students’ experiences, including whether adipose tissue was salient or not within the course, whether or not a change in perception occurred, perceived importance of adipose tissue, anatomical distribution, and personal or practical implications. Thematically similar responses were grouped into broader categories. Representative quotations were selected to illustrate each theme and to reflect the range of student perspectives within each course.

Given the exploratory and educational nature of this study, qualitative findings were used to contextualize and complement quantitative survey results rather than to generate theory.

## Results

### Quantitative results

A total of 63 students completed the survey. Of these, 50 responses were from ANAT 403 (50/173 students; 28.9% response rate) and 13 were from ANAT 503 (13/17 students; 76.5% response rate). All respondents completed all multiple-choice survey items.

### Perceptions of adipose tissue and different body sizes

Ordinal logistic regression indicated that students in ANAT 503 had significantly higher odds of endorsing more positive conceptualizations of adipose tissue along an ordered scale from harmful/pathological to vital organ system compared to students in ANAT 403 (OR = 4.24, z = 2.32, p = 0.021; Fig. 1). Through multinomial models, students in ANAT 503 were significantly more likely than students in ANAT 403 to report a positive change in attitudes toward adipose tissue (Q9; Fig. 2), rather than no change (RRR = 32.4, 95% CI: 3.48–301, p = 0.002), as well as to report a change that was different but neither explicitly positive nor negative (RRR = 37.0, 95% CI: 3.56–3.84, p = 0.003). In contrast, for attitudes toward people with different body sizes (Q10), there were no statistically significant differences between courses in the likelihood of reporting a positive change (RRR = 0.21, 95% CI: 0.17–2.52, p = 0.56), a negative change (RRR = 1.33, 95% CI: 0.29–6.02, p = 0.95), or a different but neutral change (p > 0.90). Similarly, for reported changes in feelings about one’s own body (Q11), no statistically significant differences were observed between courses for positive (RRR = 1.82, 95% CI: 0.30–10.9, p = 0.51), negative (RRR = 0.58, 95% CI: 0.00–518, p = 0.92), or different but neutral changes (RRR = 1.14, 95% CI: 0.13–1.14, p = 0.91).

**Figure 1:**
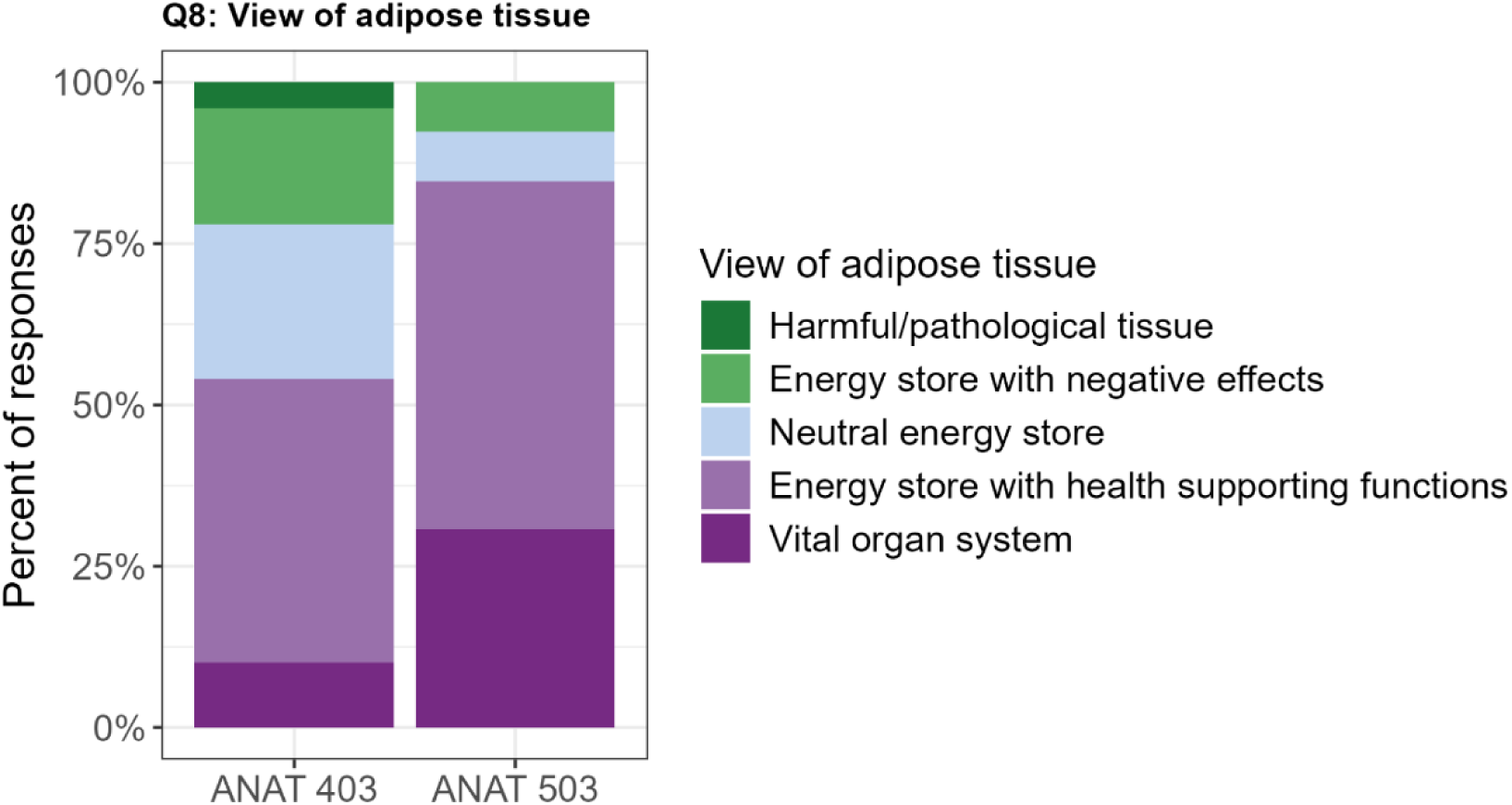
Student conceptualizations of adipose tissue in ANAT 403 and ANAT 503. ANAT 503 students were more likely to endorse adipose tissue as a vital organ system or as an energy store with health-supporting functions, whereas ANAT 403 students more commonly endorsed neutral to negative energy storage or pathological views.

**Figure 2:**
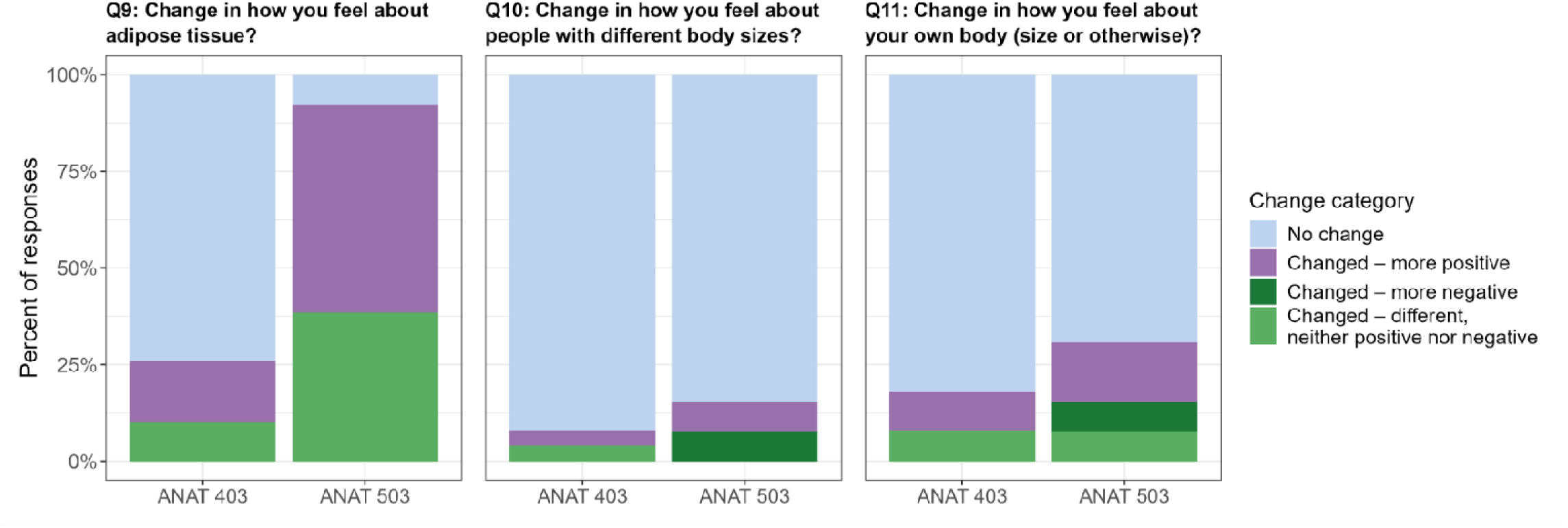
Self-reported changes in attitudes toward adipose tissue (Q9), people with different body sizes (Q10), and one’s own body (Q11) in ANAT 403 and ANAT 503. Differences between courses were most apparent for attitudes toward adipose tissue, with ANAT 503 students reporting more positive or neutral change, whereas responses to body-related questions were similar across courses.

### Validated weight bias metrics

Ordinal logistic regression revealed no statistically significant differences between courses in levels of agreement for either validated weight bias item. Students in ANAT 503 showed higher odds of stronger agreement that weight-based discrimination should be taken as seriously as other forms of discrimination (λ = 0.868), and lower odds of agreement that body weight is not a reliable indicator of health (λ = 0.568), though neither association reached statistical significance (Q13: OR = 1.84, p = 0.31; Q12: OR = 0.50, p = 0.25; Fig. 3).

**Figure 3:**
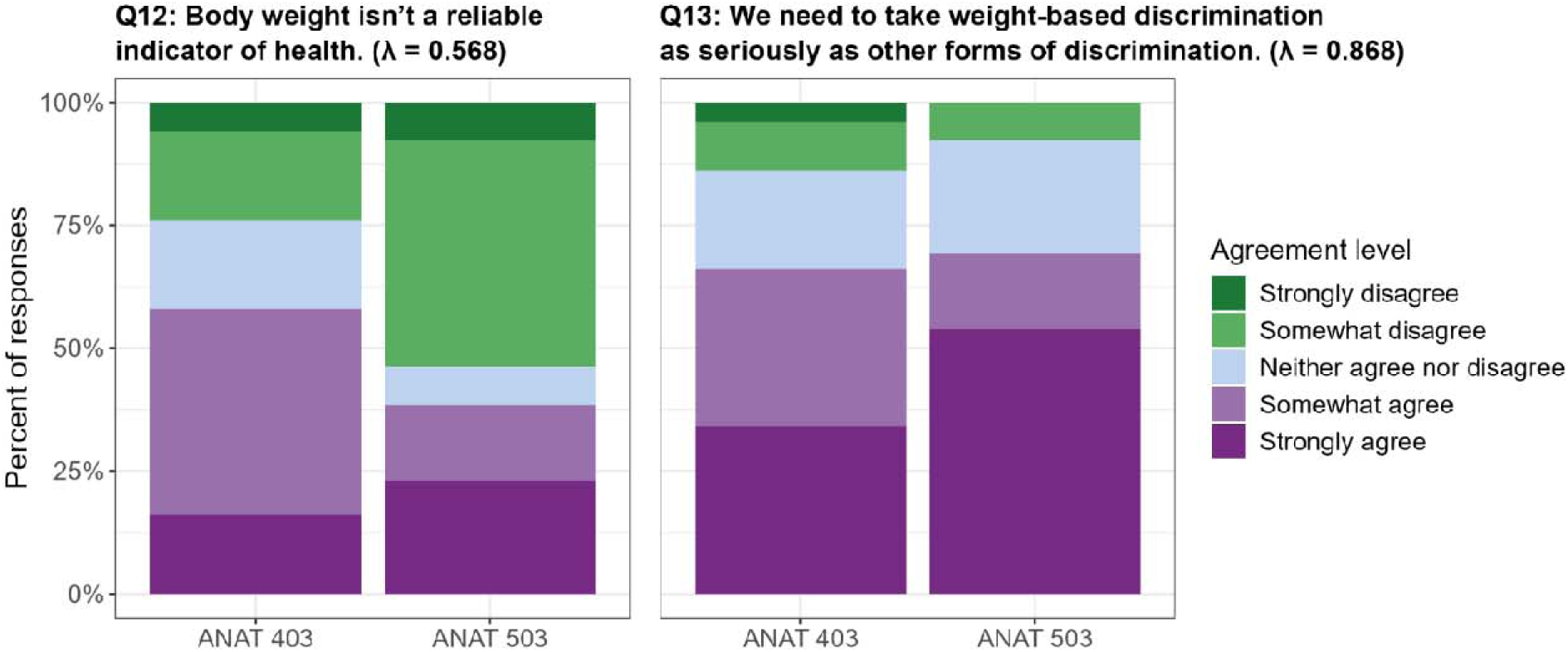
Distribution of student responses to weight-related attitude questions Q12 (“Body weight isn’t a reliable indicator of health”) and Q13 (“We need to take weight-based discrimination as seriously as other forms of discrimination”) in ANAT 403 and ANAT 503. Bars show the percentage of responses across agreement categories. λ values shown in panel titles denote factor loadings reported in the original validation study (Cain et al., 2019). Although no statistically significant differences were detected between courses for either question, ANAT 503 students showed a greater tendency to somewhat disagree with body weight as an indicator of health and to strongly agree with the seriousness of weight-based discrimination.

### Assessment of adipose-related anatomical understanding

ANAT 503 student performance on adipose-related exam questions was consistently high across the semester (Fig. 4), with mean performance exceeding 75% across most assessments. A substantial proportion of students achieved full correctness on adipose-related items at each time point. However, variability among students was evident, with some individuals demonstrating partial or no correct responses on specific assessments. Performance patterns were broadly similar across quizzes and practical examinations, and generally tracked the class’ overall assessment averages.

**Figure 4:**
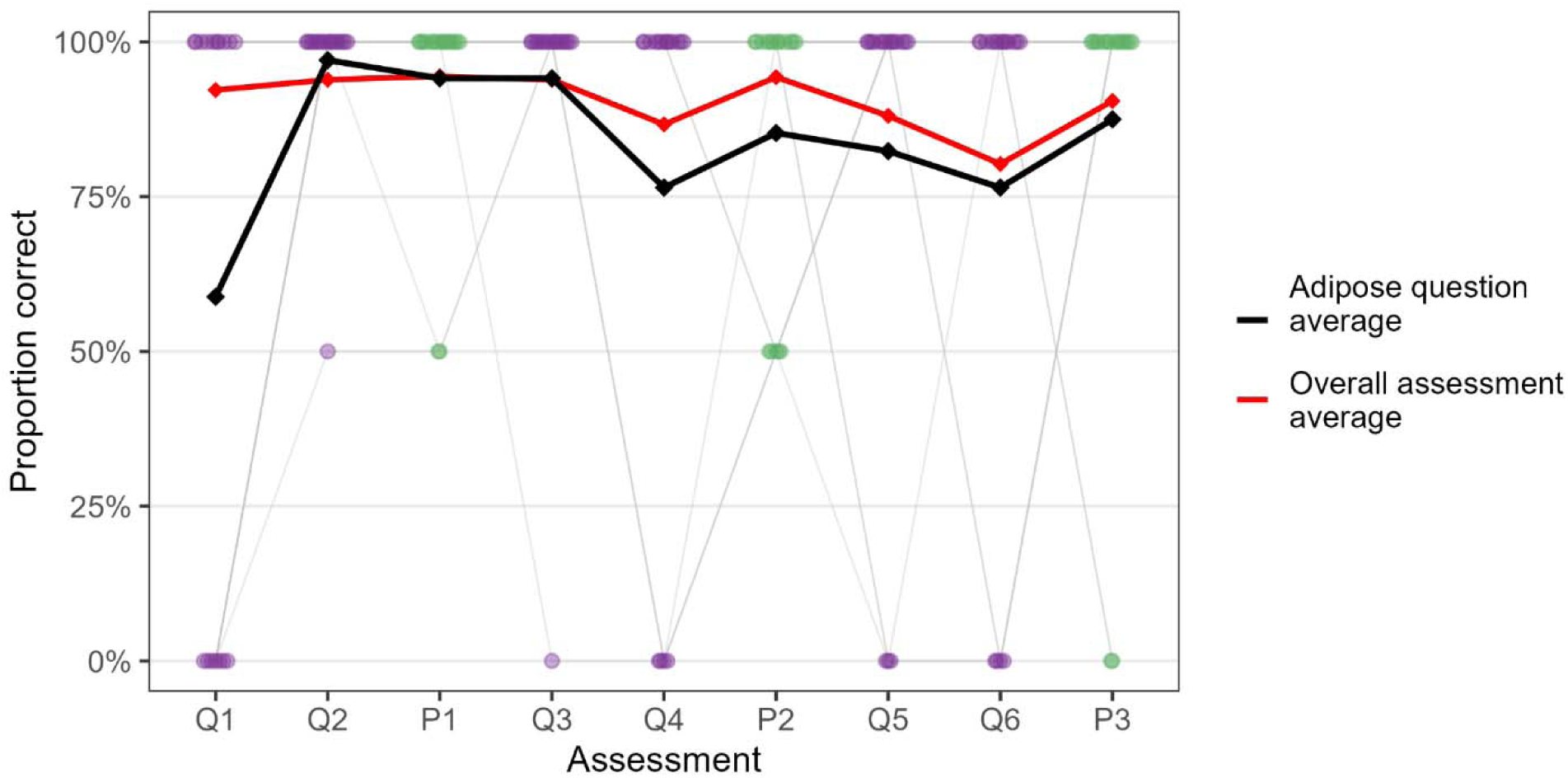
Student performance on adipose-related exam questions across the semester. Proportion of adipose-related questions answered correctly by individual students across sequential assessments (Q1–Q6: quizzes; P1–P3: practical exams). Each point represents a student’s proportion correct on adipose-related items for a given assessment, with gray lines connecting repeated measures. Points are colored by assessment type (quizzes: purple; practical exams: green). The black line represents the mean student performance on adipose-related questions, and the red line the overall class average for each assessment. Performance on adipose-related questions remained high across most assessments (>75%) and generally tracked the overall class average.

### Qualitative results

Analysis of open-ended responses revealed clear qualitative differences between students enrolled in the adipose-inclusive anatomy dissection course (ANAT 503) and those in the standard anatomy course (ANAT 403).

### Anatomy 503

Students in ANAT 503 were overall more likely to respond to the free form text boxes on questions related to adipose tissue than they were to questions related to body size. This aligns with the quantitative results of where they report more change in perception, as such, the majority of emergent themes are related to adipose tissue as opposed to body size. These themes included vitality and importance, ubiquitousness and diversity, and practical and personal concerns (Table 1).

**Table 1:** Representative free-response comments from ANAT 503 (intervention group) illustrating emergent themes related to adipose tissue, including vitality and importance, ubiquity and diversity, and practical or personal concerns. Participant gender is indicated in brackets.

| Vitality/importance | Ubiquitousness/diversity | Practical/personal concerns |
| --- | --- | --- |
| "It is critical for physical and metabolic support in all areas of anatomy."<br>"Colloquially fat is seen negatively. The lab has changed that for me, both in my perceptions scientifically (more important rather than discardable) and in real life (my fat is important!)."<br>"I think the anatomy lab has shown me how necessary fat and other viscera is to act as filler space in the body. I think it's cool to see how it holds the kidneys in place and its function as the mesentery and omentum of the abdominal organs."<br>"Became more aware of how important this layer is for protection of other structures."<br>"I never realized how many vital functions it had." | "I didn't realize how vital adipose tissue is and the amount of places it is in that we wouldn't necessarily think of, like the eyes or the hands."<br>"I never realized how pretty much every structure is supported by adipose tissue. It isn't just externally on the abdominal muscles but rather protecting vasculature, holding organs in place, etc."<br>I have seen the variety and various uses and placements it has in the human body.<br>There is so much more than I thought and it is everywhere! | "It definitely changes the dissection if there is excess amounts of fat, but is present in every single body and is essential for function."<br>"I feel like I see certain depots as negative whereas I see others as positive, for example I see the omentums positively, but I was not fond of the abdominal adipose tissue."<br>"I am more conscious of what I eat now after seeing so many dried and clogged arteries. I think also looking at an older cadaver's body made me value taking care of my body more since I want to prevent as many of the negative effects of aging as I can." |

### Vitality and importance

Students in ANAT 503 frequently described adipose tissue as a vital and functionally important component of human anatomy. Many emphasized its role in physical and metabolic support across body systems, noting that their experience in the anatomy lab enlightened them to its importance. One student wrote, “*I never realized how many vital functions it had*.” Another student reflected on a broader shift in perspective: “*Fat is seen negatively. The lab has changed that for me, both in my perceptions scientifically (more important rather than discardable) and in real life (my fat is important!*”.

### Ubiquitousness and diversity

Responses from ANAT 503 students also highlighted increased awareness of the ubiquity and diversity of adipose depots throughout the body, including depots not typically emphasized in anatomy education. One student noted, *“I never realized how pretty much every structure is supported by adipose tissue. It isn’t just externally on the abdominal muscles but rather protecting vasculature, holding organs in place, etc.*” Another similarly reflected, *“I didn’t realize how vital adipose tissue is and the amount of places it’s in that we wouldn’t necessarily think of, like the eyes or the hands*.”

### Practical and personal concerns

Some students did note technical difficulty of working with a donor with perceived higher quantities of adipose tissue. One student wrote, “*It definitely changes the dissection if there is excess amounts of fat*,” while also emphasizing that adipose tissue “*is present in every single body and is essential for function*.” Others expressed mixed feelings towards the tissue, particularly as it related to the perceived health of the donors. As one student explained, “*I feel like I see certain depots as negative whereas I see others as positive, for example I see the omentum’s positively, but I was not fond of the abdominal adipose tissue.”* A small number of responses reflected personal health concerns prompted by the laboratory experience, such as, “*I am more conscious of what I eat now after seeing so many dried and clogged arteries.”*

### Anatomy 403

Students in ANAT 403 (Table 2) were overall less likely to engage with free form text responses related to either adipose tissue or body size, and when they did respond, they most often reported that these topics were not salient components of the course. This aligns with the quantitative results indicating minimal reported change in any measured perceptions following the anatomy laboratory experience. As a result, few emergent themes were identified beyond lack of salience and lack of change in perceptions.

**Table 2:**
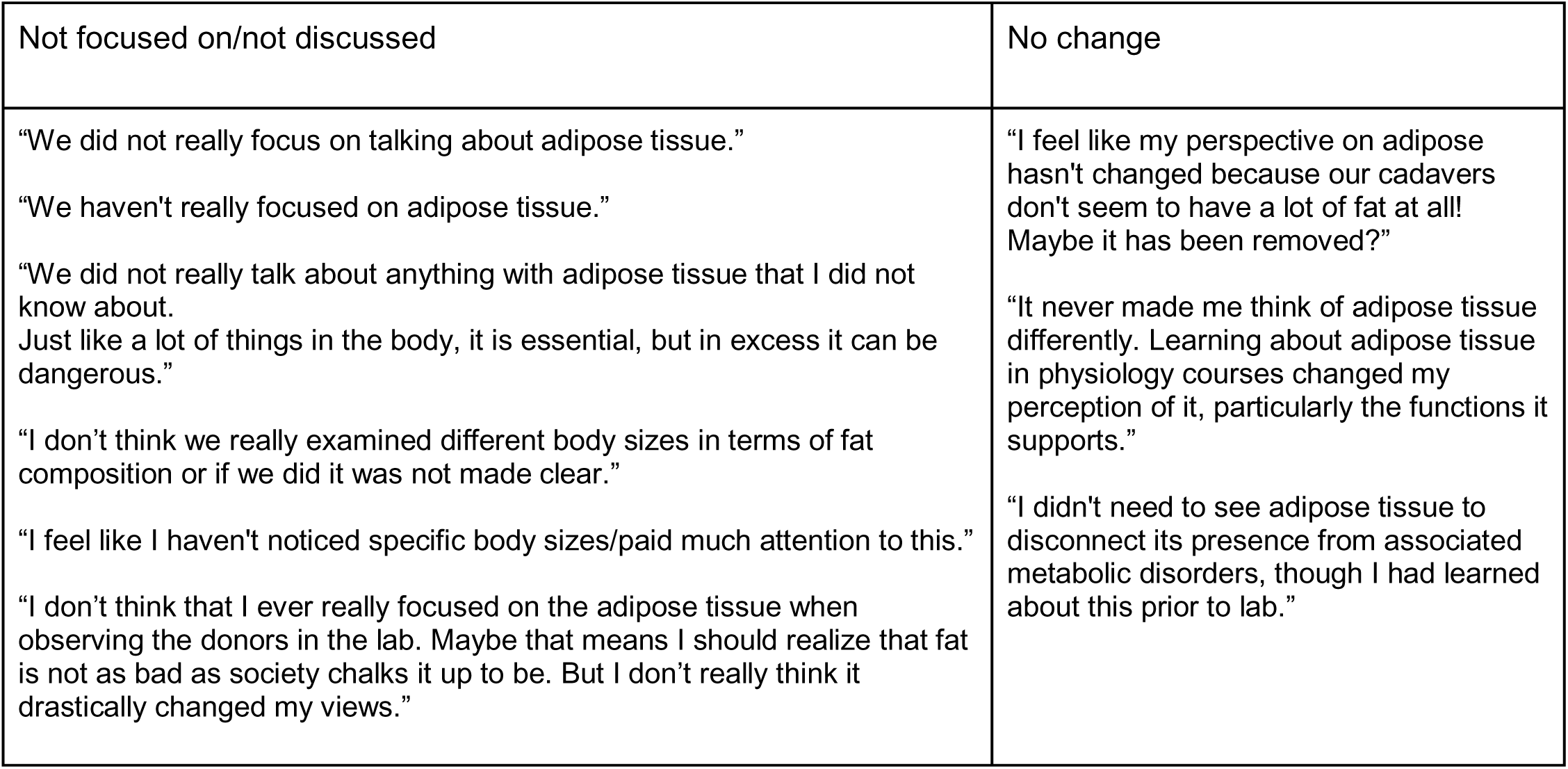
Representative free-response comments from ANAT 403 (control group) illustrating emergent themes of limited curricular emphasis on adipose tissue and self-reported lack of change in perceptions. Participant gender is indicated in brackets.

### Adipose tissue and diversity in body sizes not salient in coursework

ANAT 403 students commonly reported that adipose tissue was not a major focus of the course. As one student stated, “*We haven’t really focused on adipose tissue,*” a sentiment that was echoed across responses. These responses support that adipose tissue is typically largely absent from the anatomy curriculum. Additionally, ANAT 403 students report being unable to identify diversity in body sizes from their laboratory materials. One wrote, “*I don’t think we really examined different body sizes in terms of fat composition or if we did it was not made clear*.” Another questioned, “*Our cadavers don’t seem to have a lot of fat at all! Maybe it has been removed?”* These responses likely reflect the use of plastinated donors, where adipose tissue is typically fully removed whenever possible.

### No reported changes in perceptions

Consistent with these observations, many ANAT 403 students indicated little to no change in their perceptions of adipose tissue or body size from anatomy lab. When perspectives on adipose tissue were described, they were attributed to prior coursework in physiology rather than the anatomy laboratory experience itself. One student noted, “*It never made me think of adipose tissue differently*,” while another explained, “*Learning about adipose tissue in physiology courses changed my perception of it, particularly the functions it supports.*”

## Discussion

Our findings suggest that inclusion of adipose tissue in anatomy dissection is associated with improved conceptualizations of adipose tissue, without evidence of worsening perceptions related to body size or weight bias. Students enrolled in the adipose-inclusive dissection course (ANAT 503) were significantly more likely to describe adipose tissue as a vital organ system and to report changes in their understanding of its functional importance, whereas students in the non-dissection anatomy course (ANAT 403) reported minimal perceptual change. In addition, student performance on adipose-related exam questions was consistently high across the semester, suggesting that adipose tissue was effectively understood as an anatomical component across assessments.

The most consistent effect observed in this study was a shift in how students conceptualized adipose tissue itself. Quantitatively, students in ANAT 503 were more likely to endorse views of adipose tissue as a vital organ system and to report positive or neutral changes in their perceptions of adipose tissue. Qualitatively, these changes were reflected in themes emphasizing adipose tissue’s vitality, functional importance, and widespread anatomical presence. Students frequently described gaining a new appreciation for adipose tissue as structurally supportive, metabolically active, and integral to human anatomy.

A comparison between student reflections in the present study and those reported by Goss et al. highlights how pedagogical framing within the dissection environment may shape perceptions of adipose tissue. In Goss et al., students frequently described adipose tissue using language of invasion, disgust, and pathology, characterizing fat as “gross,” “suffocating,” or something that “invades” the body, and often extending these interpretations to moral and health judgments about people with larger bodies. In contrast, students in the adipose-inclusive dissection course more commonly used language of function, support, and integration, describing adipose tissue as “essential for function,” “protecting” and “supporting” structures throughout the body. Although some students noted technical challenges or expressed ambivalence toward specific depots, these reflections were generally contextualized within an understanding of adipose tissue as a universal and biologically necessary component of human anatomy.

In contrast to perceptions of adipose tissue, we did not observe statistically significant differences between courses in attitudes toward people with different body sizes or in validated measures of weight bias. Importantly, the comparison course (ANAT 403) here functions as a negative control for dissection-related effects, as students learn anatomy without human anatomical donor dissection. As dissection itself has been identified as a context in which negative perceptions of adipose tissue and larger bodies may emerge; we would expect students in the adipose-inclusive dissection course to report more negative attitudes than students in the non-dissection course. Instead, students in the dissection course showed similar responses on measures of body size perceptions and validated weight bias items, indicating that an anatomy dissection course with the inclusion of adipose tissue does not exacerbate negative attitudes related to body weight or size.

However, while non-significant, we observed a trend toward greater disagreement with the statement that “body weight is not a reliable indicator of health” among students in the adipose-inclusive dissection course. This finding suggests that while reframing adipose tissue as a functional organ system may improve anatomical understanding and may help to prevent worsening of negative perceptions related to body size and weight stigma, further improvements in attitudes toward body size and weight stigma will likely require additional educational efforts that directly address body diversity, health, and social biases.

These findings suggest that integrating adipose tissue into anatomy dissection curricula is both scientifically appropriate and pedagogically beneficial. Treating adipose tissue as an object of study rather than an obstacle to be removed aligns anatomy education with contemporary biomedical understanding and supports more accurate representations of human variation. Importantly, this approach does not appear to worsen student attitudes toward body size and may help counteract some of the negative perceptions previously associated with traditional dissection practices. Future work in anatomy education should build on this foundation by pairing adipose-inclusive curricula with more explicit instruction on body size diversity and health outcomes. Such efforts should also recognize that weight-related stigma is a pervasive societal phenomenon with influences far beyond the anatomy classroom.

This study has several limitations. Sample sizes were modest, particularly for the adipose-inclusive course, and response rates differed substantially between courses, which may have limited statistical power. Outcomes were assessed using self-reported survey data, which may be influenced by social desirability bias, particularly for questions related to weight bias and attitudes toward body size. While validated weight bias items were included, a full weight bias scale was not administered, and long-term changes in attitudes were not assessed. Additionally, our comparison group consisted of a non-dissection anatomy course rather than a dissection-based course without adipose-inclusive instruction, which limits our ability to isolate the effects of adipose inclusion from those of dissection itself. Future studies should address these design limitations and directly compare dissection-based courses with and without adipose-focused instruction. This work will help to clarify how improvements in anatomical understanding can be integrated into broader efforts to reduce bias in medical education.

## Conclusions

We conclude that inclusion of adipose tissues in anatomy course curricula is not only more scientifically holistic, it may be a promising first step towards improving negative perceptions related to body size and weight bias associated with anatomy dissections. However, more work is needed to continue to improve negative weight biases associated with anatomy education and in the overarching medical field, and we suggest this could be done with more targeted education on diversity in body sizes and health in addition to adipose tissue inclusion.

## Acknowledgements

We gratefully acknowledge the anatomical donors and their families. This work was supported by the University of Michigan’s Lecturers’ Employee Organization Professional Development Fund.

## Notes

### Competing Interest Statement

The authors have declared no competing interest.

